# Advancing Cardiac Graft Assessment Methods During Normothermic Machine Perfusion to Achieve Improved Transplantation Outcomes

**DOI:** 10.1101/2025.02.20.639392

**Authors:** Emmanuella O. Ajenu, Manuela Lopera Higuita, Maya Bolger-Chen, George Olverson, Khanh T. Nguyen, Gurjit Singh, Padraic Romfh, Doug Vincent, S. Alireza Rabi, Asishana A. Osho, Shannon N. Tessier

## Abstract

Heart failure, a leading global health challenge, affects over 23 million people worldwide, with heart transplantation being the gold standard for end-stage disease. However, the scarcity of viable donor hearts presents a significant barrier, with only one-third of available grafts used due to stringent selection criteria. Machine perfusion technologies, particularly normothermic machine perfusion (NMP), offer promise in improving graft preservation and assessment, yet their full potential for predicting transplantability remains underexplored. This study investigates three assessment methods to enhance human heart evaluation during NMP, focusing on mitochondrial function, left ventricular (LV) performance, and inflammatory markers. First, resonance Raman spectroscopy (RRS) is employed to assess mitochondrial redox state as a proxy for metabolic competency, offering a non-invasive and dynamic evaluation of mitochondrial function during ex vivo preservation. Second, LV function is quantified using intraventricular balloons, providing critical insights into graft viability and performance. Third, inflammatory markers and endothelial activation are assessed from perfusate to predict post-transplant outcomes. These methods were tested on human donor hearts declined for transplantation, preserved via static cold storage (SCS) and subsequently assessed with NMP in Langendorff mode. The results demonstrate that these parameters can be easily integrated into existing clinical perfusion workflows and hold potential for improving heart transplantation outcomes by enhancing graft selection and optimizing donor heart use. Future studies will further validate these biomarkers across different preservation techniques and evaluate their clinical applicability.

## Introduction

Heart failure is a major healthcare issue, impacting over 23 million individuals globally, with 6 million affected in the United States alone [1]. The progressive and incurable nature of this condition results in heart transplantation being the gold standard of care for end-stage disease, as it increases quality of life and life expectancy [2]. However, despite the remarkable advances in surgical techniques, immunosuppressive therapies, and post-operative care, the scarcity of viable hearts remains a critical barrier [3]. The global increase of heart failure has heightened the demand for heart transplantation, while the supply of donor grafts has remained stagnant [4–6]. This stagnation is due to various factors, including logistical challenges in organ procurement, allocation, and transportation, as well as ethical considerations surrounding consent and organ donation [7]. Along with these, extremely stringent selection criteria based on current assessment methods lead to consequentially low donor graft usage (only 1 of every 3) [8–11]. In effect, more sensitive monitoring of heart quality will ensure improved identification of suitable grafts leading to increased utilization of extended-criteria donors [12].

Many techniques, such as temperature-controlled static cold storage, hypothermic machine perfusion, and normothermic machine perfusion (NMP), have been developed over the years to address the heart shortage crisis [13, 14]. Despite some of these techniques offering many advantages such as transport duration, preservation of metabolic status, and/or simplicity, only normothermic machine perfusion (NMP) using the Organ Care System (OCS) by Transmedics has been FDA approved for the transportation/assessment of hearts donated after circulatory death (DCD), a subset of organs previously deemed unsuitable for transplantation [15, 16].

Despite these important achievements and advances, the field has yet to exploit the full potential of predicting graft transplantability during normothermic machine perfusion.

In recognition of this potential, significant effort has been dedicated to advancing assessment methods during NMP in order to improve the evaluation of donor heart viability. NMP maintains the heart in a near-physiological state allowing the incorporation of various assessment metrics, such as perfusion variables, metabolic and injury biomarkers, hemodynamic values and functional metrics [17]. Although the predictive capacity of some of these metrics have demonstrated mixed results (i.e. lactate profiles), others have shown promise especially in combination (i.e. oxygen uptake rate in combination with contractile function metrics), highlighting the potential of dual assessment approaches [18]. Other metabolic biomarkers, including high energy phosphates, have indicated preservation of metabolic status is central for optimal graft preservation [19], while some functional metrics have been reported to have a strong correlation with post-transplantation outcomes [20, 21]. Building on these efforts [22–24], the aim of this study was to explore three assessment methods that could be easily incorporated into the existing workflow of diverse perfusion systems, and that offers a comprehensive evaluation of multiple variables, including metabolic status, function, endothelial activation and inflammatory markers.

The first assessment method evaluates the redox state of the mitochondria as a proxy for the graft’s metabolic competency post-storage [25]. Mitochondria are indispensable for metabolic homeostasis and function [26], and, in particular, their redox state is indicative of important functional features. For example, under oxygen limiting conditions, either during cold or warm ischemia, the number of reduced mitochondria may increase, a potential indicator of the degree of ischemia that could be correlated with heightened injury and loss of function. Further, upon the re-introduction of oxygen during MP, a shift towards a reduced state may indicate insufficient oxidative phosphorylation and dysfunctional mitochondria, while an overly oxidized state may suggest increased oxidative stress that exacerbates bioenergetic insufficiencies [27–29]. Despite the promise of the redox state to define mitochondrial function, current techniques to evaluate mitochondria are either non-specific or necessitate destructive tissue processing, limiting its potential during *ex vivo* graft assessment prior to transplantation. Instead, our group has introduced the use of resonance Raman Spectroscopy (RRS) to measure the redox state of mitochondria dynamically from the surface of *ex vivo* grafts [23, 24]. Importantly, this device enables these measurements to be made without the need for tissue sampling and can be performed throughout the preservation and assessment phases irrespective of temperature. Using a high-resolution spectrometer and 441 nm excitation wavelength, unique RRS fingerprints of the oxidized and reduced forms of mitochondrial cytochromes are quantified as the ratio of reduced to total mitochondria (or Resonance Raman Reduced Mitochondrial Ratio: 3RMR) [22].

The second assessment method determines left ventricular (LV) function using intraventricular balloons. Retention of LV function is of paramount importance for the success of heart transplantation due to its crucial role in maintaining adequate cardiac output and overall cardiovascular performance. Marginal hearts may exhibit varying degrees of dysfunction or damage, underscoring the importance of LV function assessment to determine graft transplantability. Importantly, clinically used NMP systems perfuse in Langendorff mode without ventricular filling. Hence, intraventricular balloons would be relatively easy to incorporate into existing system designs and have been widely used experimentally to assess LV function [30–32]. Typically, a pressure transducer connected to a saline-filled balloon is inserted into the left ventricle. This enables the quantification of left ventricular pressure, as well as contractility and relaxation parameters, which have been shown to correlate with cardiac loading capacity [33].

The third assessment method involves the quantification of endothelial activation and inflammatory markers. Inflammatory cytokines have long been associated with graft dysfunction, with levels of tumor necrosis factor-α (TNF-α) shown to predict right ventricular failure early after transplantation [34] and elevated levels of, both, TNF-α and interlukin-6 measured in cardiac grafts deemed non-transplantable when compared to transplantable grafts [35]. Expectedly, the concentration of these inflammatory markers along with various others is utilized in donor evaluation processes due to their prognostic information [36]. Similarly, endothelial cell activation has been pre-clinically associated with reduced posttransplant outcomes [12]. As access to circulating perfusate during NMP is readily available, the quantification of endothelial activation (e.g. p-selectin, sVCAM-1, sICAM-1, ADAMTS13), pro-and anti-inflammatory cytokines (IL-8, IL-10, MCP1, TNFɑ) from perfusate, instead of biopsies, provides the ability to non-invasively implement their prognostic capabilities into any perfusion system.

This manuscript demonstrates the potential of three assessment parameters taken together with human hearts that were declined for transplantation, including the mitochondria redox state, left ventricular contractility and relaxation, and inflammatory markers. These assessment parameters are proposed (in part) due to ease of integration into existing clinical perfusion workflows, which perfuse in Langendorff mode. Moreover, we choose a study design whereby human hearts were transported in static cold storage (SCS), followed by onsite normothermic machine perfusion in Langendorff mode. This enables us to separate preservation from assessment, thereby opening the potential for techniques proposed herein to be applied irrespective of the choice of preservation method, especially with respect to the mitochondrial redox state. While we choose to demonstrate proof-of-principle with SCS, future studies will further evaluate other preservation techniques, including hypothermic machine perfusion or continuous normothermic machine perfusion, as well as validate the clinical utility of these biomarkers with transplanted hearts.

## Methods

### Organ acquisition

This study was conducted in accordance with approval from the institution review board (IRB). Human hearts rejected for transplantation due to suboptimal donor criteria were retrieved by the New England Donor Services (NEDS, Waltham, MA, USA) and transported under hypothermic storage in ice-cold storage solution. Donor and graft characteristics are shown in Table 1.

| Donor ID | A | B | C | D | E | F | G | H | Summary |
| --- | --- | --- | --- | --- | --- | --- | --- | --- | --- |
| Age(years) | 66 | 20 | 40 | 47 | 44 | 63 | 45 | 38 | 45.3(13.5) |
| Sex(M/F) | M | M | M | F | M | F | M | F | 63% (M) |
| WIT (mins) | 16 | 24 | 0 | 0 | 0 | 0 | 0 | 0 | 5(8.8) |
| CIT (mins) | 60 | 180 | 240 | 420 | 294 | 300 | 339 | 270 | 262.8(100.8) |
| Death Type | DCD | DCD | DBD | DBD | DBD | DBD | DBD | DBD | DCD(n=2)<br>DBD(n=6) |
| LVEF(%) | No Echo | 55 | 50 | 63 | 70 | 55-60 | 60 | 40-45 | 59.6(6.8) |
| Additional notes | CAD | Moderate tricuspid regurgitation | Trace mitral/tricuspid regurgitation | 65mins cardiac arrest | Mild mitral/trace tricuspid regurgitation | Trace mitral/tricuspid regurgitation | Mild mitral/trace tricuspid regurgitation | Trace mitral/mild tricuspid regurgitation<br>81 mins cardiac arrest |  |
| WIT, Warm Ischemic Time; CIT, Cold Ischemic Time; DCD, Donation by Circulatory Death; DBD, Donation by Brain Death; LVEF, Left Ventricular Ejection Fraction; CAD, coronary artery disease; Mean(STDev) |  |  |  |  |  |  |  |  |  |

### Normothermic Machine Perfusion (NMP)

All chemicals were purchased from Sigma Aldrich unless otherwise stated. All hearts were perfused in Langendorff for 6 h in an organ chamber (VentriFlo, Pelham, NH USA) at 37°C with 4L of perfusate containing 88.98 mM sodium chloride, 3.35 mM potassium chloride, 4.59 mM calcium chloride dihydrate, 0.51 mM sodium dihydrogen phosphate dihydrate, 15 mM sodium bicarbonate, 1.18 mM magnesium dichloride hexahydrate, 10.95 mM glucose monohydrate, 0.125 mM dextran 40, 2.28 mM bovine serum albumin, heparin (2000 U/L), insulin (200 U/L), 0.4% penicillin-streptomycin, and hydrocortisone (200 U/L). After perfusate was brought to volume by the addition of distilled water, sodium hydroxide (5 M) was added to bring pH to 7.8, followed by the addition of HEPES for pH stabilization which brought the pH of the solution to 7.4.

The details of the perfusion circuit are discussed in detail elsewhere [37]. Briefly, perfusate was added to the circuit and circulated by a pulsatile pump (VentriFlo, Pelham, NH USA) through a pediatric oxygenator (Medtronic, Minneapolis, MN) flushed with 100% O_2_. A Windkessel bag (VentriFlo, Pelham, NH USA) was implemented before the aorta to provide a more physiological source of coronary perfusion [37]. After arrival, hearts were weighted, instrumented and connected to the perfusion system while continuously monitoring aortic flow and pressure via flow (Transonic, Knoxville, TN) and pressure (iWorx, Dover, NH) sensors, as previously described [37]. The initial 500 mL of perfusate was collected and discarded to remove high potassium originating from the storage solution. Rhythmic compressions were done on the hearts to avoid ventricular distention until intrinsic contractions were observed. In case of fibrillation, hearts were defibrillated (Hewlett Packard, Louisville, KY) starting with 10 J, with subsequent shocks administered in increments of 10 J until rhythmicity was observed. A back up pacer was set at 86 beats per minute.

### Viability assessment

*Perfusate blood gas analysis*: Oxygen, lactate, pH and potassium measurements were obtained from perfusate samples via blood gas analysis (Siemens Medical Solutions, Malvern, PA, USA) 20 mins after initiating perfusion and every half an hour thereafter. Oxygen uptake rate (OUR) was determined following the formula below:

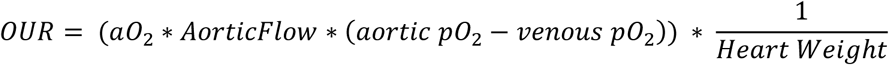

Where aO_2_ is oxygen solubility coefficient (0.00314 µLO_2_/mmHgO_2_/mL); Aortic pO_2_ is the partial oxygen pressure (mmHg) of perfusate going to the coronaries; Venous pO_2_ is partial oxygen pressure (mmHg) of perfusate coming out of the coronary sinus; Aortic Flow is flow rate (ml/min); and heart weight is graft weight (g) [38].

*Perfusion hemodynamics, weight gain, and tissue sampling*: Vascular resistance was calculated by dividing the coronary pressure by flow rate and normalized for weight of the heart before the initiation of perfusion. Hearts were weighed before and after perfusion, and these values were utilized to calculate percent weight gain, a proxy for edema. Biopsy punches (5 mm) were harvested from the full thickness of the left ventricle before initiation of perfusion (pre – samples), individual samples were flash frozen or stored in 5% formaldehyde. After the 6 h of the perfusion, a circumferential section was fixed in 5% formaldehyde and larger sections of the heart were flash frozen in liquid nitrogen for downstream analysis. All samples in formaldehyde were fixed for a minimum of 24 h and transferred to 70% Ethanol.

*Microscopy*: Pre – and post – perfusion samples, both from the full thickness of the left ventricle were embedded in paraffin and subsequently stained with H&E and TUNEL-labeled fluorescein-labeled dUTP (Roche, Basel, Switzerland), as described previously [39]. All images were acquired with NanoZoomer (Hamamatsu Photonic, Shizuoka, Japan). Four images from each TUNEL-stained slide were acquired at 20x in semi-random locations and utilized for quantification of percent cell death via MATLAB (MATLAB®, The MathWorks, Inc., Natick, MA).

*Perfusate-based pro-inflammatory markers*: Pro-inflammatory markers were measured via bead-based immunoassay (Luminex, Austin, TX), by Eve Technologies Corp (Alberta, CA). Briefly, outflow perfusate samples at 20 mins (T0.5) and 6 h (T6) were collected, frozen and shipped on ice for processing. Analysis was performed using a Luminex 200 system, following the kit’s manufacturer instructions (MilliporeSigma, Burlington, MA).

*Quantification of mitochondria redox state via Raman Resonance Spectroscopy (RRS)*: Mitochondria redox state was quantified via a portable Raman Resonance Spectroscopy (Pendar Technologies, Cambridge, MA). The device uses a 441 nm excitation laser and 9 mW power source coupled to a compact probe head (dimensions are 8 mm by 12 mm by 60 mm) that was fixed at 5-8 mm above the left ventricle for a 2 mm spot measurement, as shown in Supplementary Fig. 1A. RR photons were collected via a temperature-controlled detector over the course of a 180 s measurement time. The spectral readout was analyzed using custom LabView software, whereby the fluorescence baseline was subtracted from raw spectrum and the remaining signal was the convolved RR spectrum. To demultiplex the combined spectrum (also see [22]), the employed regression algorithm determined the optimal ratios of chromophore spectra from a previously recorded library to fully explain the measured spectrum. The library components for this analysis include mito-reduced, mito-oxidized and myoglobin, as shown in Supplemental Fig. 1B. The regression utilizes a large region of the spectrum (700-1700 cm^-1^), and small differences in each chromophore spectra are accounted for in the results. In addition, the high resolution of the spectrometer allows the distinction of very small shifts in the position of related peaks, as shown in Supplementary Fig. 1C. This regression determines the relative concentration of metabolic molecules (i.e. mitochondria, myoglobin) by calculating the coefficient in the equation (Supplemental Fig. 1C) as a weighted sum of each component’s spectrum. The resulting coefficients are utilized to calculate the ratio between reduced mitochondria and total mitochondria (sum of reduced and oxidized), this metric is denoted as 3RMR.

*Functional assessment*: Left ventricular pressures were acquired via a latex balloon attached to a fluid-filled silicon tube and pressure sensor (iWorx, Dover, NH). Briefly, after 45 minutes of perfusion time, the deflated balloon was inserted through the left atrium, and advanced through the mitral valve to reach the left ventricle. Once situated, the balloon was filled with saline until pressure during diastole was 40 mmHg. The left ventricular pressure was recorded for 15 mins every hour using Labscribe software (iWorx, Dover, NH, USA), and the data processed with MATLAB (The MathWorks Inc, Natick, MA). Contractility and relaxation metrics were obtained by calculating the maximum and minimum derivative, respectively, of the LV pulse pressure waves [37].

## Statistical Analysis

To determine the relationship between pre-perfusion 3RMR values and function, Pearson’s correlation coefficients were computed via Prism (GraphPad Software Inc., La Jolla, CA). Pearson’s r was used to assess the strength and direction of linear relationships between the variables. Data represented as median ± interquartile range.

## Results

### Conventional assessment metrics, alone, are unreliable to determine graft viability

Viability markers are denoted conventional when they can be obtained from un-modified Langendorff perfusion from analyzing perfusate samples (Lactate, pH, potassium, oxygen uptake rate) or through common perfusion parameters (pressures/flows – vascular resistance). Perfusate lactate levels are broadly utilized, both in clinic and laboratory settings, as one of the few quantitative markers of cardiac graft health during NMP [40–46]. Clinical operating procedures recommend the use of lactate levels as a proxy for proper perfusion. Both a downward trend in total lactate, as well as confirmation of lactate consumption by the hearts (i.e. concentration of outflow lactate is lower than the concentration of inflow lactate) are broadly used to determine favorable metabolic signatures of ex vivo perfused hearts [47].

Most of the hearts utilized for this study seemed to alternate between consuming (ΔLactate _outFlow–inFlow_ < 0) and producing (ΔLactate _outFlow–inFlow_ > 0) small amounts of lactate resulting in ΔLactate _venous–aortic_ oscillating around zero for most of the perfusion time (Fig. 1A) with a couple exceptions. Hearts A and G demonstrated a downward trend of ΔLactate _outFlow–inFlow_ for most of the perfusion time, indicating lactate consumption. Heart F experienced large fluctuations in consumption and production resulting in large shifts in ΔLactate _outFlow–inFlow_. Despite most hearts maintaining ΔLactate _outFlow–inFlow_ close to zero, lactate in perfusate increased over time in most hearts (Fig. 1B). The accumulation of lactate in the perfusate is accompanied by a decrease in pH over time which resulted in perfusates with pH as low as 6.6 (heart C) by the end of the perfusion time (Fig. 1C). Unsurprisingly, however, lactate is not solely responsible for pH changes, as the pH in heart G still dropped (-0.15) even when accumulating no lactate over the entire time of perfusion. Alternatively, no changes in pH were observed in the perfusate of heart B despite a slight accumulation of lactate.

**Figure 1:**
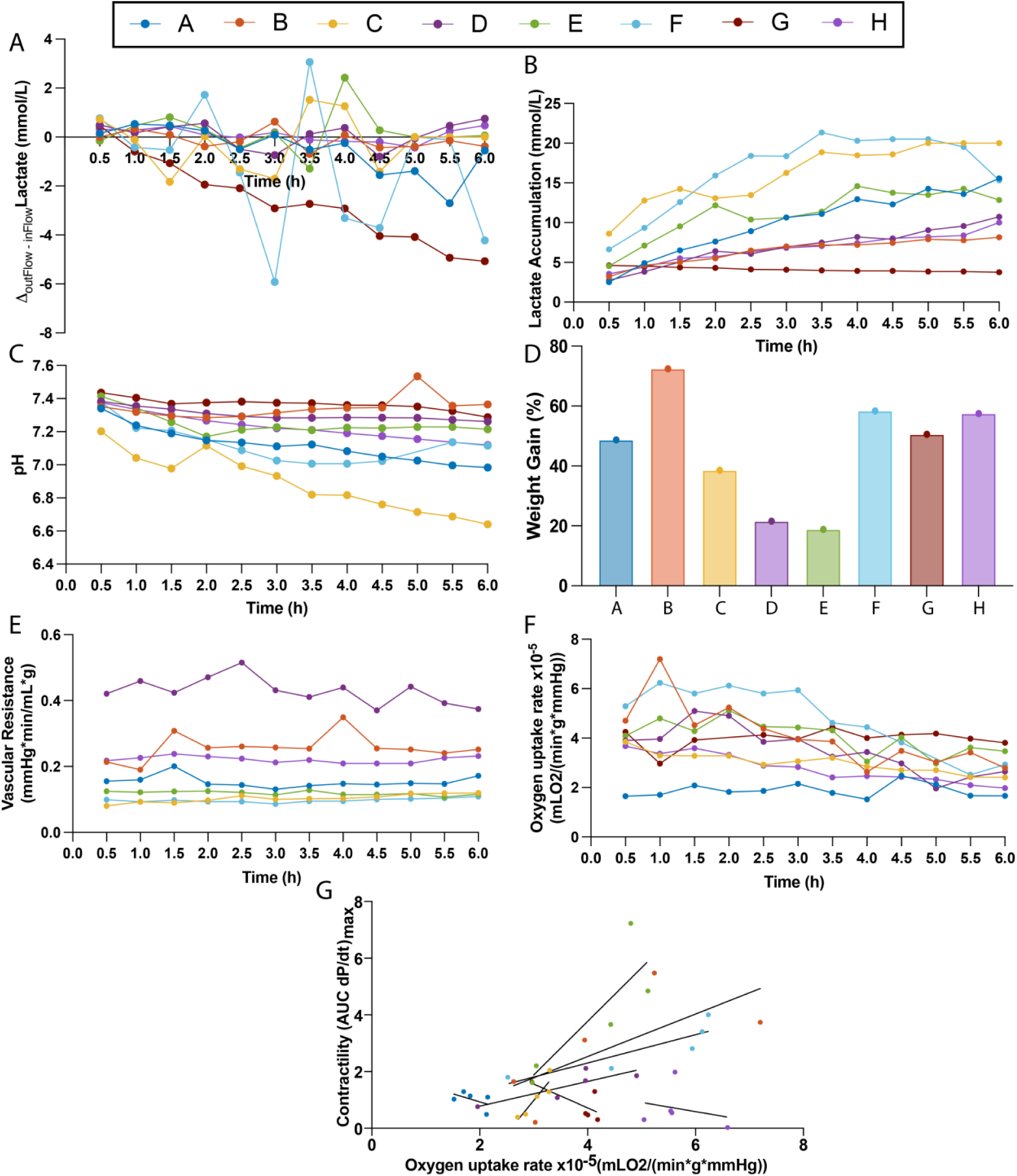
Conventional assessment metrics of graft viability. Metrics are considered conventional if their acquisition is possible during un-modified Langendorff perfusion. (A) Lactate consumption, defined as the difference between inflow and outflow lactate, overtime during NMP. (B) Lactate accumulation in perfusate measured overtime from inflow perfusate. (C) Perfusate pH over perfusion time. (D) Percent weight gained during NMP, a proxy for cardiac edema. (E) Vascular resistance. (F) Oxygen uptake rate. (G) Correlation between oxygen consumption and cardiac contractility (AUC dP/dt_max_).

Regardless of the amount of lactate accumulated or changes in pH, all hearts in this study experienced major weight gain, indicating substantial edema (Fig. 1D). This edema appears more prominent in the right ventricular wall of most cardiac grafts than in the left ventricular wall, as demonstrated by H&E histological images, likely due to the decreased resistance posed by the thinner wall of the right ventricle (Fig. 2). Despite differing magnitudes of vascular resistance between hearts – i.e. heart D (0.427 ± 0.0581 mmHg*min/(mL*g)) vs. heart F (0.096 ± 0.008 mmHg*min/(mL*g)) – small to no changes were seen for any of the hearts over time (Fig. 1F). Alternatively, a slight downward trend in oxygen consumption (oxygen uptake rate) is seen in all hearts with the exception of heart A (Fig. 1G). All three hearts with comparatively better initial cardiac function (Hearts B, E and F, see Fig. 3) and two of the less functional hearts (Heart B and D) have a positive correlation between oxygen consumption and cardiac contractility, indicating a decrease in cardiac contractility in response to decreased oxygen consumption (Fig. 1G). Whereas three out of the four hearts with initially low cardiac function (Hearts A, G, H) demonstrated a negative correlation between cardiac contractility and oxygen consumption.

**Figure 2:**
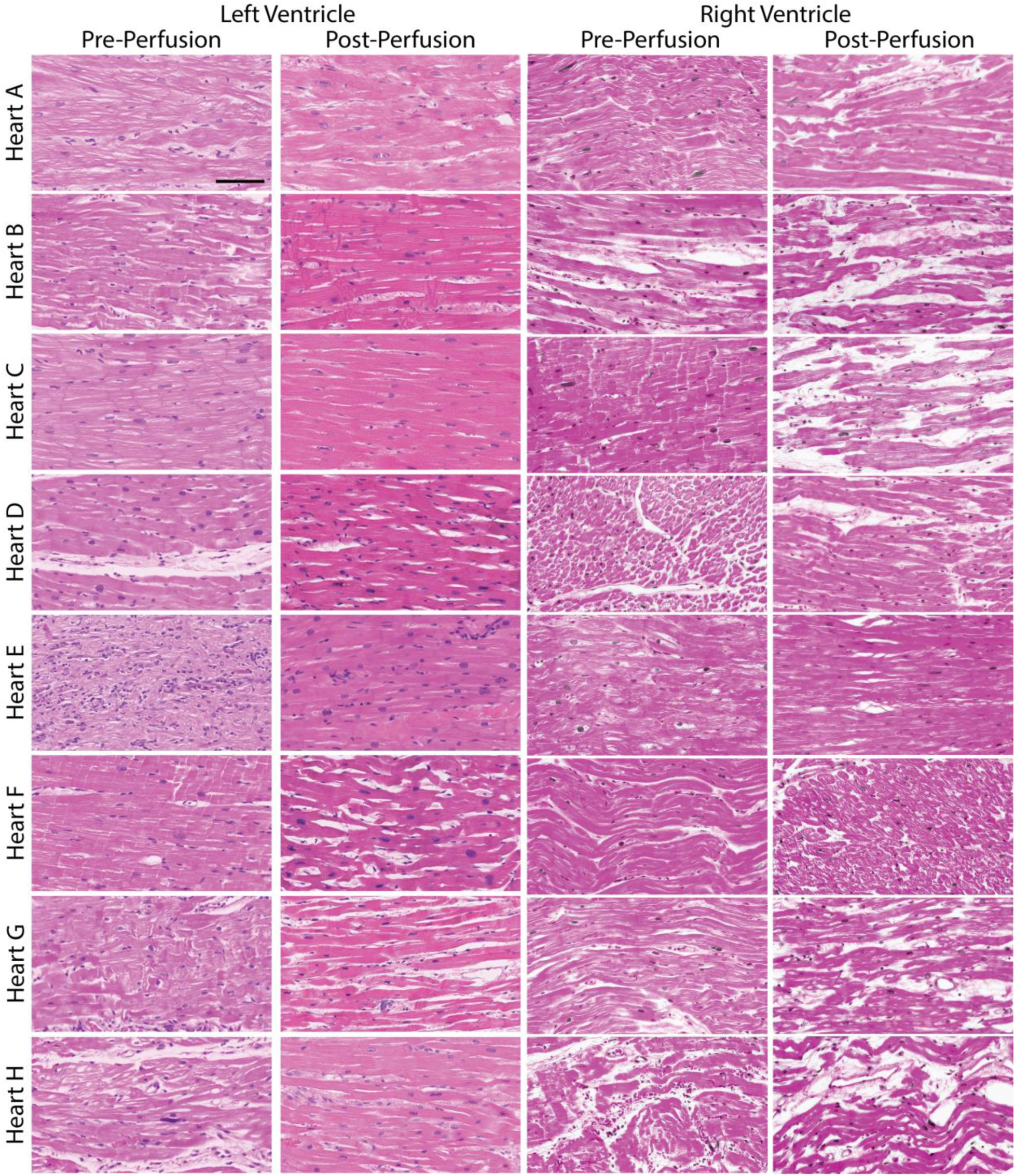
Histological Analysis of Cardiac Grafts Pre-and Post-NMP. (A-H) Representative Hematoxylin & Eosin (H&E) staining images of cardiac grafts sampled from the left and right ventricular wall at two time points: pre-perfusion, following transport and storage, and post-perfusion, after 6 hours of NMP. Scale bar 100 μm.

**Figure 3:**
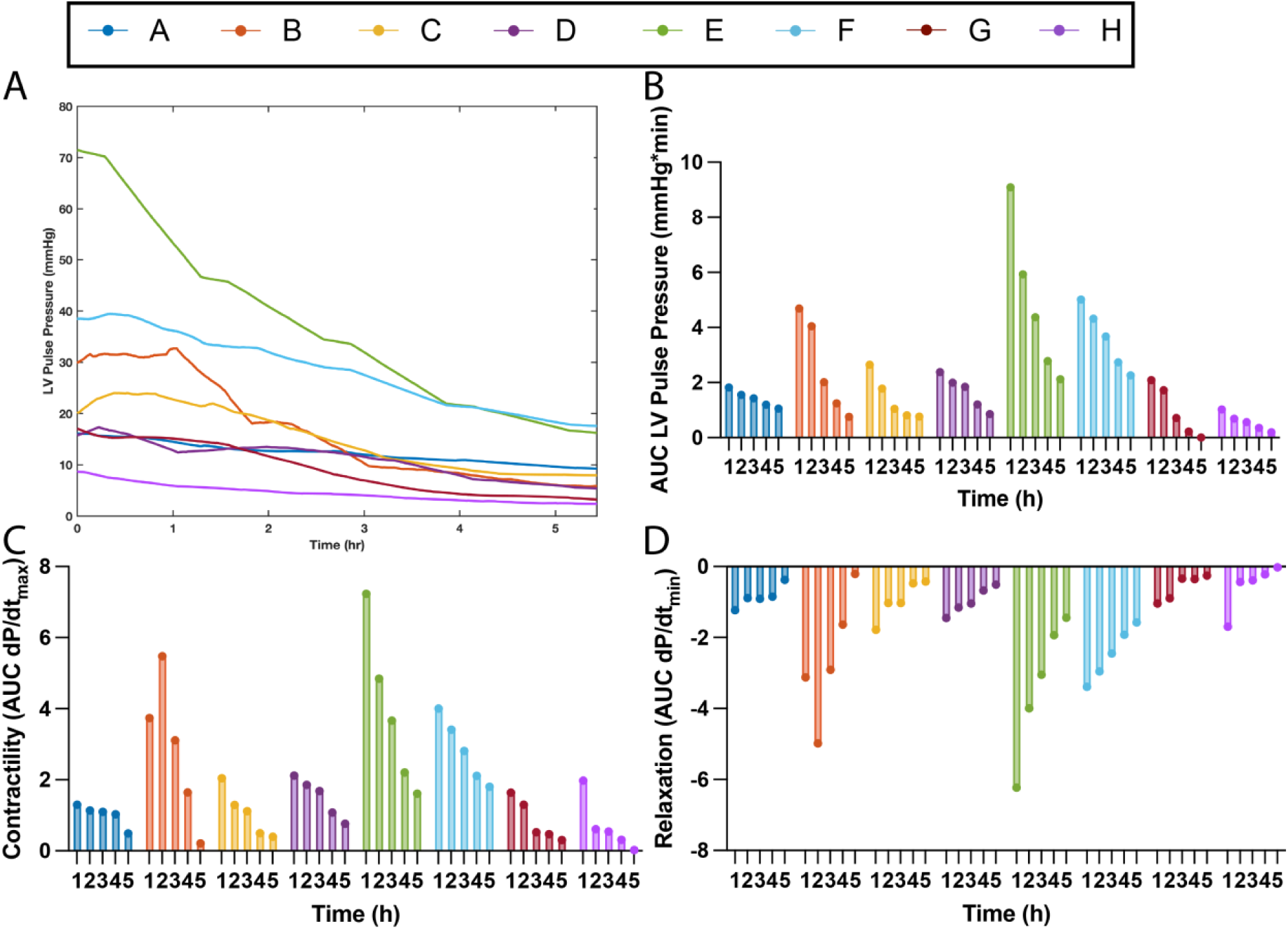
Nonconventional assessment metrics – left ventricular pulse pressure. Metrics are considered non-conventional if their acquisition is not common practice clinically. (A) Maximum systolic pressure plotted over time, denoted as left ventricular pulse pressure (LVPP). (B) The area under the curve (AUC) of the LVPP trend for every hour of perfusion. (C) Cardiac muscle contractility quantified from the maximum derivative of the pressure pulse. (D) Cardiac muscle relaxation quantified from the minimum derivative of the pressure pulse.

### Left ventricular (LV) function improves graft assessment during machine perfusion in Langendorff mode

Most hearts perfused in this study demonstrated similar and difficult-to-differentiate biochemical patterns during perfusion (lactate, pH, O2 consumption, etc.). The exclusive use of these parameters to evaluate cardiac grafts in clinical settings highlights the importance of other easily implemented evaluation techniques, which are further investigated in this study. For instance, the use of intraventricular balloons to assess ventricular function. Since intraventricular balloons do not require ventricular filling, they are relatively easy to incorporate into existing clinical system designs that perfuse in Langendorff mode. Through LV balloon placement, it was determined heart E had the highest initial LV pulse pressure outputting 71 mmHg, followed by heart F which outputted 38 mmHg (Fig. 3A & 3B). Interestingly, despite initiating perfusion at very different LV pulse pressure magnitudes, the pulse pressure for both hearts ended at similar magnitudes after 6 h of perfusion (**E:** 16 mmHg, **F**: 17 mmHg). This implies heart E experienced a steeper loss in function, while heart F experienced a more gradual loss of function. Heart B initiated perfusion with a LV pulse pressure around 30 mmHg and was able to maintain this moderate function for an hour before the LV pulse pressure dropped, ultimately reaching ∼5 mmHg by the end of perfusion. The remaining hearts barely recovered ventricular function resulting in initial pulse pressures being between 8 – 20 mmHg and final LV pulse pressures 2 – 10 mmHg. Irrespective of initial LV pulse pressure, all hearts experienced a steady decline in LV pulse pressure throughout perfusion, suggesting severe LV dysfunction and lack of transplantability. Furthermore, the decrease in LV pulse pressure was accompanied by a decrease in both, contractility (Fig. 3C) and relaxation (Fig. 3D) over time in all hearts, with the exception of Heart B. This heart, despite the loss LV pulse pressure, experienced an increase in contractility (**1h**: 3.77 vs. **2h:** 5.47) and relaxation (**1h**:-3.12 vs. **2h:-**4.98) when comparing the first hour of perfusion to the second.

### Mitochondrial redox state may aid graft assessment during static cold storage and reanimation

The redox state can be quantified by cross referencing the mitochondrial cytochromes’ unique resonance Raman spectral signatures, when excited at specific wavelengths, with spectral libraries. This process allows for the calculation of the ratio of reduced to total mitochondria (Resonance Raman Reduced Mitochondrial Ratio – 3RMR). Since the main role of mitochondrial cytochromes is to carry electrons across the ETC to oxygen, a balance between oxidized and reduced cytochromes is indicative of a healthy metabolic state, which is defined as a 3RMR between ∼10-20, suggesting efficient energy production [22, 48]. Alternatively, higher numbers of reduced cytochromes (i.e. high 3RMR) can indicate ischemia stress, while over-oxidized mitochondria (i.e. low 3RMR) can indicate reperfusion injury as damaged cells no longer consume oxygen.

In accordance with this, hearts displaying lower 3RMR values (21% ± 2, Fig. 4A) before the initiation of perfusion (on-ice measurements, Hearts E & F) exhibited higher left ventricular pulse pressures (Fig. 3A & 3B), contractility (Fig. 3C), and relaxation (Fig. 3D) in the initial hour of perfusion compared to hearts with higher on-ice 3RMR (74% ± 30.5, Hearts A, C, D & G). These results demonstrate a strong correlation (r^2^ = 0.725) between on-ice 3RMR values and cardiac contractility in the initial hour of perfusion. The exception to this trend was Heart B, which displayed completely reduced mitochondria in the on-ice Raman measurement (100% 3RMR) but produced slightly higher cardiac work than hearts A, C, D & G.

**Figure 4:**
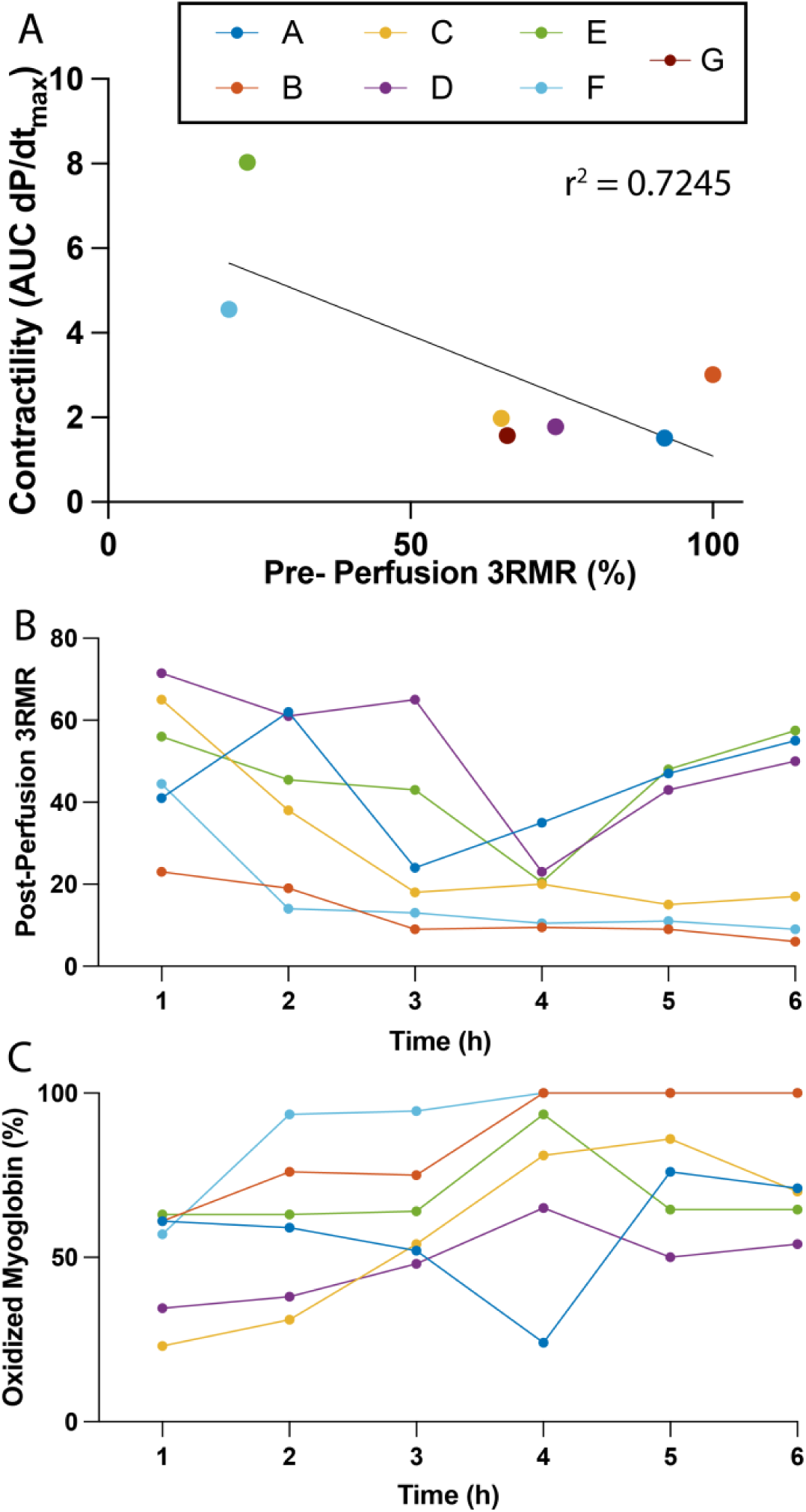
Nonconventional assessment metrics – Mitochondria redox state via Raman resonance spectrometry. (A) Correlation between pre – perfusion 3RMR values and cardiac contractility in the first hour of perfusion. (B) Over time 3RMR values during cardiac perfusion. (D) Percent oxidized myoglobin over perfusion time.

### All marginal cardiac grafts were deemed unsuitable for transplant based on 3RMR

Generally, the results in this manuscript indicate these marginal grafts were not transplantable, despite most cold ischemia times being within clinically acceptable range and warm ischemia times from 0-24 mins (Table 1). For instance, none of the hearts were able to reach optimal mitochondria function determined by balanced ratios of reduced and oxidized mitochondria (i.e. mid-range 3RMR values, Fig. 4B). Instead, all grafts demonstrate 3RMR values in unhealthy extremes with grafts B, C, and F reaching an abnormally low 3RMR values after the first couple hours of perfusion, while grafts A, E & D maintained abnormally high 3RMR values throughout the experiment, indicating an unhealthy accumulation of reduced mitochondria. Furthermore, based on myoglobin levels, a known oxygen reservoir, a progressive decline in myocyte oxygen utilization was observed in most grafts during perfusion time, indicating a gradual loss of mitochondrial function (Fig. 4C). The initial slightly lower percentage of oxidized myoglobin at 1 h suggests some degree of oxygen delivery from myoglobin to myocytes, indicative of mitochondria respiration via the ETC. The over-time increase in the ratio/percentage of oxidized myoglobin indicates an accumulation of oxygen in these molecules, suggesting the inability of myocytes to efficiently utilize the oxygen. Despite some degree of function in the initial hour of perfusion, the post-storage suboptimal conditions of these grafts were supported by hematoxylin & Eosin (H&E) staining that revealed pre-perfusion cardiomyocytes lacked essential structural elements such as striations and intercalated discs, indicating sub-optimal tissue integrity in many of the hearts (Fig. 2).

### Normothermic machine perfusion is highly pro-inflammatory

Increased inflammatory activation in cardiac grafts is associated with higher percentages of immunorecognition, rejection and other cardiopathies in transplant patients [49, 50]. As a result, analysis of inflammatory and adhesion biomarkers can provide valuable insight into the viability of cardiac grafts, either, when preserved or assessed via NMP. Despite the heterogenicity of the fold increase magnitude, all hearts utilized in this study demonstrate a large increase in secretion of key pro-inflammatory molecules (Fig. 5). Adhesion molecules such as soluble intercellular adhesion molecule-1 (sICAM-1, Fig. 5A) and soluble vascular cell adhesion molecule-1 (sVCAM-1, Fig. 5B) were found to be slightly elevated to different degrees across all grafts, likely indicating endothelial cell activation. Interestingly, the increased secretion of P-selectin (Fig. 5C) and the metalloproteinase Adamts13 (Fig. 5D) was relatively uniform across all grafts, suggesting a more uniform reaction to global ischemia and reperfusion injury, consistent with its role in endothelial interactions during perfusion [51]. Similarly, the secretion of pro-inflammatory cytokines was also significant increased with the fold increase of tumor necrosis factor-alpha (TNF-α, Fig, 5D) and interleukin-6 (IL-6, Fig. 5F) being thousands of times higher at the end of the perfusion when compared with initial values. Similarly, although not as exaggerated, increased release of interleukin-8 (Fig. 5G), interleukin-1RA (Fig. 5H) and MCP1 (Fig. 5I) was also observed across all hearts in this study. Interestingly, the fold increase of anti-inflammatory cytokine interleukin-10 was also determined to be thousand times higher at the end of perfusion when compared to initial values for all hearts (Fig. 5J).

**Figure 5:**
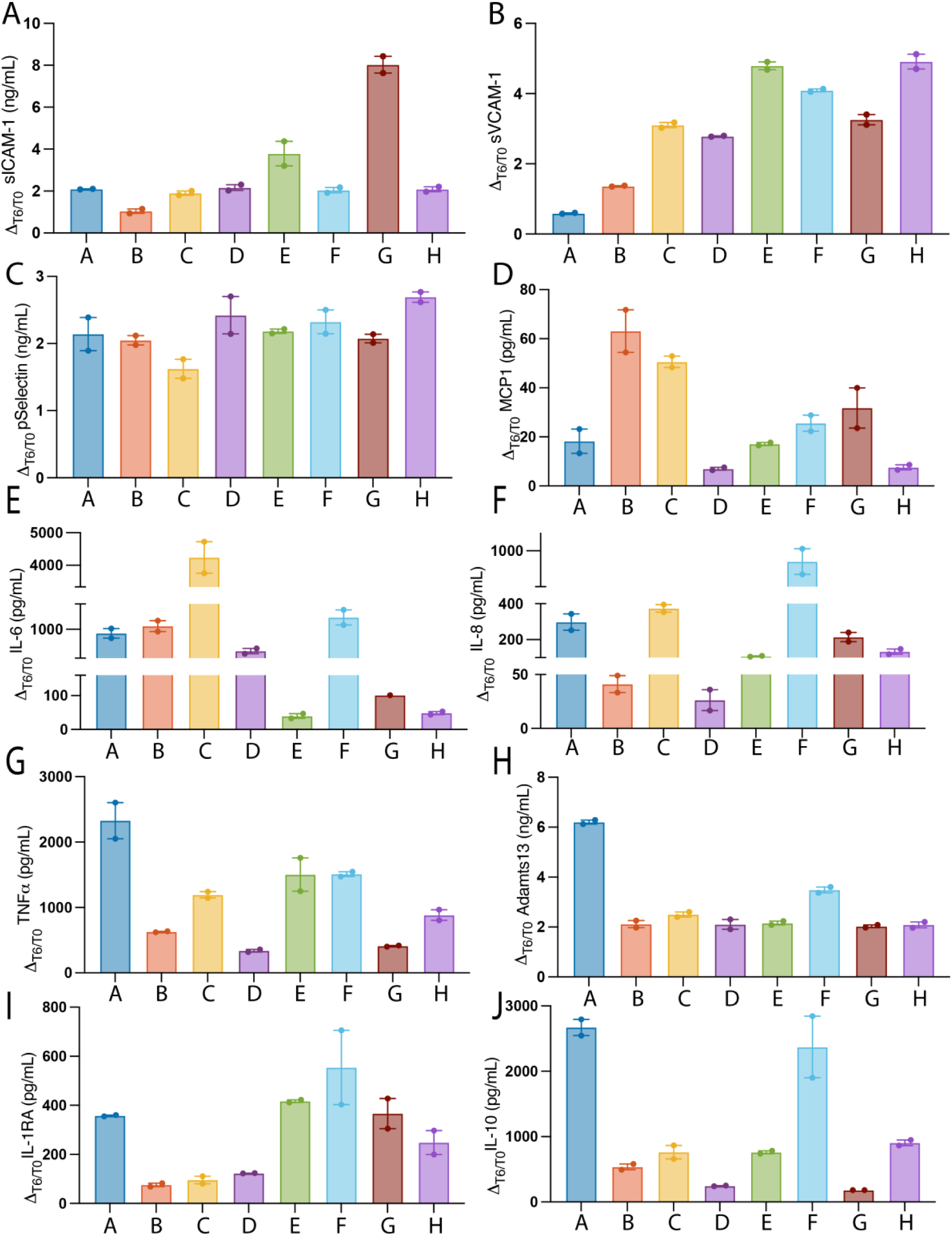
Assessment of pro – and anti-inflammatory markers secreted during normothermic machine perfusion. (A-C) In-perfusate concentration of pro-inflammatory adhesion molecules, (D) chemokine, (E-F) pro-inflammatory cytokines, pro-inflammatory chemical messenger, (H) anti-inflammatory metalloproteinase Adamts13 and (H-J) anti-inflammatory cytokines (fold increase – T_6_/T_0.5_).

## Discussion

The organ shortage crisis creates a severe mismatch between the number of patients in need and the available donor hearts, significantly hindering the power of heart transplantation as a lifesaving procedure [52, 53]. As a result, many patients face longer wait times, which lead to worsening health, increased mortality rates while on the wait list, and difficult ethical dilemmas regarding prioritization and allocation [54, 55]. One potential solution to mitigate this shortage is the use of marginal hearts – those donated after circulatory death (DCD) or by extended-criteria donors. While these organs may not meet the traditional criteria for transplantation, advances in medical technology, preservation methods (e.g., ex vivo machine perfusion), and improved surgical techniques have resulted in a slight increase in the utilization of marginal hearts for transplantation [56, 57].

Further expanding the use of marginal hearts for transplantation is the most promising strategy to address the critical shortage. However, the only way to increase the chances of successfully utilizing these organs is by identifying more definitive biomarkers that can accurately predict graft quality and transplantability. As effective, reliable methods for viability assessment do not currently exist, the aim of this study was to explore the potential of three assessment methods— resonance Raman spectroscopy (RRS) to assess mitochondrial redox state, intraventricular balloon placement for left ventricular function, and inflammatory markers from perfusate samples—to assess human grafts. Through these assessment methods, the health status of eight human cardiac grafts considered marginal was determined, both, immediately after SCS and during normothermic perfusion. It is important to note that, due to the inherent heterogeneity of these grafts, including variability in donor age, comorbidities, ischemic time, both warm and cold, implemented cardioplegia, and storage solution, which introduced significant differences in graft quality and response to preservation and reperfusion protocols, drawing definitive conclusions proved challenging. Despite these difficulties, this discussion aims to highlight interesting findings, while exploring potential causes for variability in graft outcomes and the implications for improving preservation and reperfusion strategies.

Although there is a lack of abundant corroborating clinical studies, a small number of articles suggest marginal hearts are particularly vulnerable to injury during SCS. These studies highlight that recipients of marginal hearts preserved via SCS have worse overall clinical outcomes compared to recipients of marginal hearts preserved, either by ex vivo normothermic perfusion or in Paragonix SherpaPak cardiac transport system [58–60]. The results in this study seem to support this finding with marginal hearts analyzed in this study undergoing a considerable amount of functional loss after storage, as well as important morphological changes to their cellular structures (Fig. 2, pre-perfusion). Despite five out of six DBD hearts demonstrating normal left ventricular ejection fraction in pre-procurement assessments (50%–70%, Table 1) and four out of five undergoing storage times within the clinically acceptable range, none of the hearts were able to produce peak-systolic pressures (Fig. 3A) near those considered physiological normal (90–140 mmHg) [61, 62]. This functional decline could be attributed to the known variability in cooling and increased risk of freezing injury during SCS [60, 63, 64], which has been associated with irreversible suppression of diastolic function, damage to the cardiac conduction system and denaturing of cellular proteins [65–68]. Through the implementation of RRS, information regarding the role of mitochondria redox state as an additional source of graft injury/recovery during SCS were uncovered.

Measurements of the mitochondrial redox state (reduced vs. oxidized) during the end of static cold storage seem to play a crucial role in determining the initial functional capacity of grafts after reanimation with NMP. A higher number of oxidized mitochondria indicates an availability of oxygen to act as an electron acceptor within the tissue (i.e. relatively less ischemia), likely enabling continued, albeit slow, ATP production and the maintenance of cellular homeostasis during storage. As a result, grafts containing higher number of oxidized mitochondria (or lower 3RMR) at the end of SCS showed improved graft recovery upon reanimation, as shown in Fig. 3A. In contrast, grafts with predominantly reduced mitochondria (or higher 3RMR), whereby oxygen was likely unavailable to act as an electron acceptor, showed reduced graft recovery upon reanimation. Interestingly, the depletion of oxygen reserves during SCS do not seem to be directly proportional to storage time, as some DBD grafts stored for ‘shorter’ periods of time (e.g., C & H) had a higher percentage of reduced mitochondria than DBD grafts stored for longer periods (e.g., E & F), suggesting graft to graft variability that may be related to differences in oxygen demands during SCS or may be influenced other donor characteristics. The ability of RRS to predict functional outcomes post-storage gains further importance as clinical interest in implementing hypothermic preservation techniques for marginal heart preservation rises [69].

The use of RRS also revealed warm ischemia has a significant impact on the mitochondrial redox state, even more so than prolonged cold ischemia, with both of the DCD grafts analyzed in this study (A & B) exhibiting the highest percent of reduced mitochondria when compared to all DBD hearts, despite undergoing comparatively shorter periods of SCS (60 and 180 mins, respectively). Interestingly, despite having 100% reduced mitochondria, the heart from the youngest donor (20 years, Heart B) recovered slightly more function than hearts with marginally less reduced mitochondria (A, C, D, and G, Fig. 4A). These findings may suggest that hearts from ‘older’ donors were more affected by the more reduced mitochondria, than the younger heart under similar or even less favorable conditions (due to its DCD state). The idea of a decreased tolerance of’older’ donor hearts to ischemia and reperfusion injury has been previously reported [70–73].

Age-related decline in mitochondria function has been associated with decreased production of ATP in normoxia, implying ATP levels pre-storage would have been lower in older donors, experiencing the effects of ATP depletion faster than young donor hearts [74, 75]. Furthermore, a plethora of scientific findings point to other various physiological processes crucial for ameliorating IRI becoming less effective with age. These include decreased ability to regulate N6-methyladenosine (m6A), an enzyme associated with disruptions in calcium regulation [76, 77], decreased ionic balance [78, 79], reduced autophagy and, consequently, impaired cellular quality control [80], and decreased tolerance to oxidative stress [81]. Therefore, it’s possible that better-preserved cellular mechanisms in the younger heart accounted for its ability to withstand the adverse effects of higher number of reduced mitochondria and recover slightly more function than other hearts in the cohort. Importantly, the fact that most of these grafts demonstrated normal or near-normal function pre-procurement highlights the need to augment and tailor preservation techniques to better fit the condition of the organ, as has been done for DCD hearts with normothermic preservation.

We also aimed to ascertain the cumulative effects of SCS followed by reperfusion during NMP. A lack of oxygen, in combination with slowed-down metabolism during SCS, cause mitochondria to both deplete (i.e. ATP, GTP, Coenzyme A – CoA) and accumulate (i.e. lactate, succinate) key metabolic substrates, triggering an injurious chain of reactions after oxygen is reintroduced by reperfusion during NMP (ischemia reperfusion injury – IRI) [82]. IRI is characterized by the onset of oxidative stress, inflammation, and the generation of reactive oxygen species (ROS) and reactive nitrogen species [82]. These reactive molecules damage cellular structures, including the mitochondria, leading to increased permeability and loss of integrity in the electron transport chain subunits, which further exacerbates ROS production and causes bioenergetic insufficiencies [27, 28, 83, 84]. This leads to energetic deficits, heightened cellular death and largely non-functional grafts.

In addition to significantly affecting metabolic processes, IRI also triggers a complex inflammatory response that results in transcriptional and translational upregulation of numerous pro-inflammatory cytokines (IL-6 and IL-8), glycoproteins (siCAM-1), cell adhesion proteins (sVCAM-1, p-Selectin), chemical messengers (TNF-α) and chemokines (MCP-1) [51, 85–90]. This response is further exacerbated by the pro-inflammatory effects of the non-physiological elements of normothermic machine perfusion [91]. In the case of organ transplantation, this group of pro-inflammatory mediators targets neutrophils, causing them to adhere to the vascular endothelium, leading to the obstruction of capillary beds and inhibiting complete reperfusion [92, 93]. Even in the absence of neutrophils, as in the acellular perfusions in this study, these mediators remain highly damaging. Their accumulation, particularly of TNF-α and IL-1, elicits the release of ROS, which leads to myocardial cell damage and cardiac dysfunction [92, 93]. Interestingly, due to the dual nature of inflammatory responses [94], the heightened secretion of pro-inflammatory mediators stimulated the production of anti-inflammatory cytokines (IL-1RA and IL-10), as well as the anti-inflammatory metalloproteinase ADAMTS13, all of which have been associated with protective effects during IRI [95–98]. As cytokines regulate both inflammatory and immune responses that can affect graft function and modulate vulnerability to rejection [99], the identification of the type and quantity of circulating cytokines prior to transplantation can serve as an metric for predicting transplant success.

The relevance of mitochondria function may also have value to assess reperfusion injury, as most grafts, with the exception of heart G, presented significant production of lactate during normothermic perfusion. Increased lactate production is generally caused by impaired tissue oxygenation, either from decreased oxygen delivery or a defect in mitochondrial oxygen consumption [100]. Lactic acidosis in grafts with high percentage of reduced mitochondria prior to perfusion can be attributed to mitochondria dysfunction, a known consequence of oxidative stress resulting from ROS formation and IRI [101]. However, in grafts with oxidized mitochondria prior to perfusion, their ability to produce some cardiac work for over 1 hour of perfusion suggest that the effects of IRI on mitochondrial function were less severe. Instead, lactic acidosis may be caused by decreased oxygen delivery, a consequence of high perfusate viscosity and lack of oxygen carriers.

High perfusate viscosity resulted from the implementation of high oncotic agent (bovine serum albumin) percentage, which had been reported to work well as an edema-preventive measure in other predominantly myocyte-composed organs [102]. The inverse proportionality of the oxygen diffusion coefficient to solution viscosity, along with the acellular nature of the perfusate, likely lead to limited availability of oxygen [103]. Although this oxygen unavailability likely affected all grafts in this study, the higher demand for oxygen in grafts with some degree of cardiac function resulted in lactic acidosis being induced earlier in perfusion, as demonstrated by a steeper increase in lactate accumulation within the first two hours of perfusion (Fig. 1B). This lactate accumulation led to the progressive acidification of the perfusate, which in turn diminishes calcium delivery to the cardiomyocytes, ultimately, resulting in decreased contractility and cessation of function [104]. Furthermore, fluctuations between overly oxidized and overly reduced mitochondria during perfusion (Fig. 3B) indicate that all grafts in this study were unable to achieve a mitochondrial redox state balance conducive to cardiac homeostasis, with high numbers of reduced mitochondria indicating diminished oxidative phosphorylation and metabolic activity, and overly oxidized state suggesting increased oxidative stress and mitochondrial damage.

Regardless of the source of mitochondrial dysfunction, the resultant cellular hypoxia, associated metabolic acidosis, and high lactate content were likely responsible for the significant cardiac edema observed in these grafts. Hypoxia leads to failure of membrane ATP-dependent ionic pumps due to ATP depletion, which elicits a loss of volume control within the cells [105], while metabolic acidosis and increased lactate content increase the permeability of endothelial cells [106–108]. As a result, the use of high oncotic agents in the perfusate had the opposite effect of what was intended, leading to a multifactorial onset of edema. The negative effects of high perfusate viscosity could likely be avoided by utilizing in-perfusate oxygen carriers; however, the heterogeneity of the cardiac grafts analyzed in this study made them a suboptimal model system for protocol optimization. Further research into the ideal perfusate composition, along with other protocol parameters, should be conducted in more controlled settings.

## Conclusion

The ongoing global shortage of donor hearts presents a significant challenge to heart transplantation, highlighting the need for more effective methods of graft assessment to maximize the utilization of available organs, including marginal hearts. This study explored three assessment techniques—resonance Raman spectroscopy (RRS) for mitochondrial redox state, intraventricular balloon measurements for left ventricular function, and the quantification of inflammatory markers in perfusate—to evaluate the viability of marginal human heart grafts. Despite the challenges posed by organ variability and methodological limitations, these techniques present valuable tools for refining heart preservation strategies and improving transplant outcomes. Moving forward, further studies are needed to optimize preservation protocols, validate the clinical utility of these biomarkers, and enhance graft selection to ensure the success of heart transplantation, particularly in the context of marginal and extended-criteria donor hearts. By addressing the critical gaps in heart graft evaluation, these efforts have the potential to alleviate the organ shortage crisis, improve patient outcomes, and ultimately expand the pool of transplantable hearts.

## Acknowledgements

This work was supported by generous funding to S.N.T. from the US National Institutes of Health (K99/R00 HL1431149; R01HL157803). We also gratefully acknowledge funding from the US National Institute of Health (R01DK134590; R24OD034189), National Science Foundation (EEC 1941543), American Heart Association (18CDA34110049), Harvard Medical School Eleanor and Miles Shore Fellowship, Polsky Family Foundation, the Claflin Distinguished Scholar Award on behalf of the MGH Executive Committee on Research, and Shriners Children’s Boston (Grant #BOS-85115). Further, we acknowledge the salary support to G.S provided by the Canadian Institutes of Health Research. We also thankfully acknowledge support for S.A.R Polsky Family Foundation. The authors would also like to thank the Center for Comparative Medicine and the Knight Surgical Research Lab at Massachusetts General Hospital for animal management. Finally, we thank the Mass Spectroscopy, Genomics and Proteomics, and Morphology facilities at Shriners Children’s Boston. Most importantly, we send our deepest appreciation to the donor families who have generously and selflessly enabled research with human donor organs. Finally, we extend our gratitude to our collaboration with New England Donor Services (NEDS).

---

**Supplemental Figure 1:**
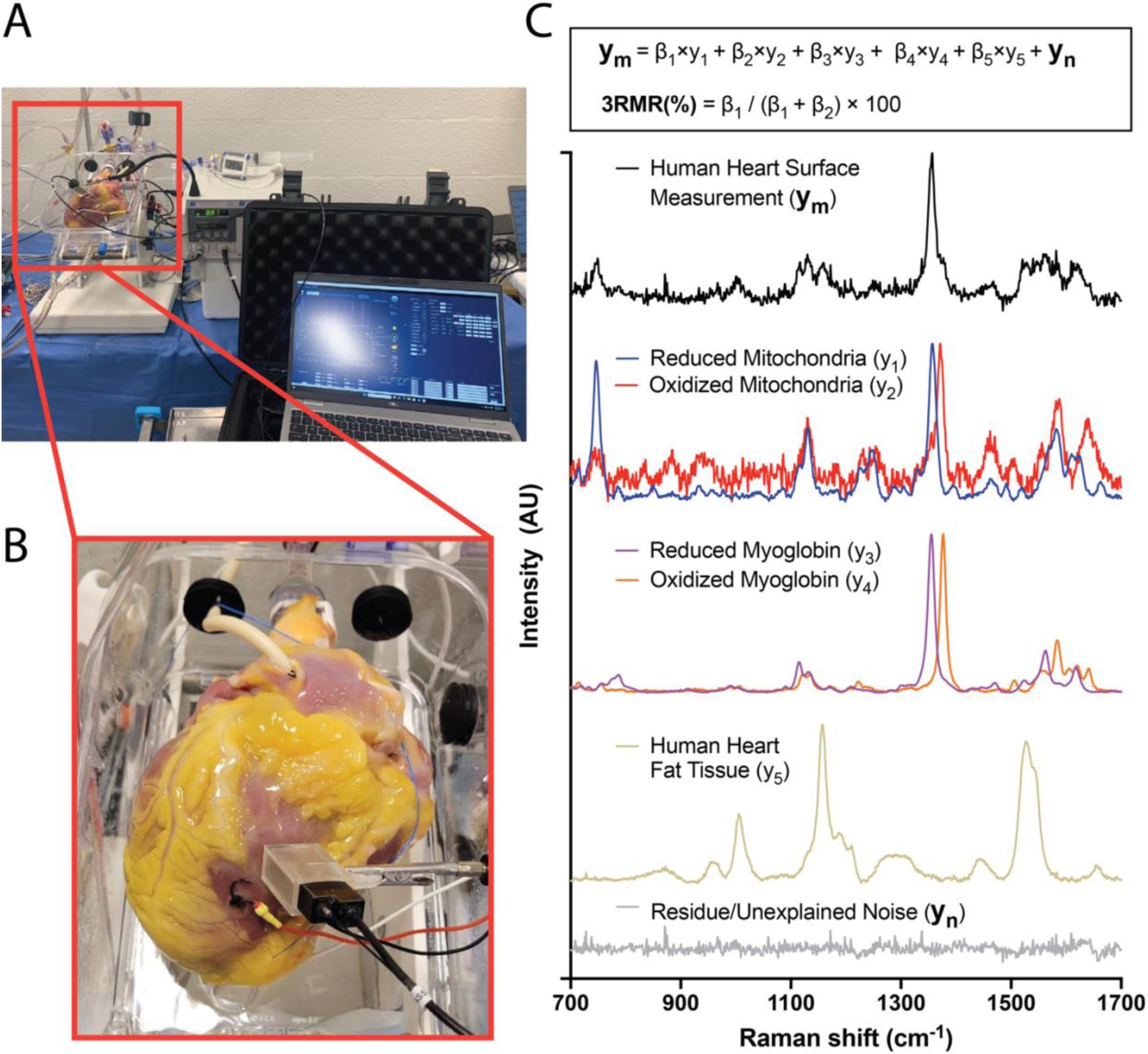
Raman Resonance Spectrometry (RRS) device setup and computational algorithm for quantification of human heart mitochondrial redox states. (A) Normothermic machine perfusion setup with portable RRS device (B) Visual of compact probe from RRS device pointed at the left ventricular surface of a human heart for non-contact measurement. (C) RRS computational regression algorithm that enables spectroscopic quantification of individual components including reduced and oxidized mitochondria, reduced and oxidized myoglobin.

